# Integrated Omics Analysis Reveals Sirtuin Signaling is Central to Hepatic Response to a High Fructose Diet

**DOI:** 10.1101/2021.09.02.458361

**Authors:** Laura A. Cox, Jeannie Chan, Prahlad Rao, Zeeshan Hamid, Jeremy P. Glenn, Avinash Jadhav, Vivek Das, Genesio M. Karere, Ellen Quillen, Kylie Kavanagh, Michael Olivier

## Abstract

**Background:** Dietary high fructose (HFr) is a known metabolic disruptor contributing to development of obesity and diabetes in Western societies. Initial molecular changes from exposure to HFr on liver metabolism may be essential to understand the perturbations leading to insulin resistance and abnormalities in lipid and carbohydrate metabolism. We studied vervet monkeys (*Clorocebus aethiops sabaeus*) fed a HFr (n=5) or chow diet (n=5) for 6 weeks, and obtained clinical measures of liver function, blood insulin, cholesterol and triglycerides. In addition, we performed untargeted global transcriptomics, proteomics, and metabolomics analyses on liver biopsies to determine the molecular impact of a HFr diet on coordinated pathways and networks that differed by diet.

**Results:** We show that integration of omics data sets improved statistical significance for some pathways and networks, and decreased significance for others, suggesting that multiple omics datasets enhance confidence in relevant pathway and network identification. Specifically, we found that sirtuin signaling and a peroxisome proliferator activated receptor alpha (PPARA) regulatory network were significantly altered in hepatic response to HFr. Integration of metabolomics and miRNAs data further strengthened our findings.

**Conclusions:** Our integrated analysis of three types of omics data with pathway and regulatory network analysis demonstrates the usefulness of this approach for discovery of molecular networks central to a biological response. In addition, metabolites aspartic acid and docosahexaenoic acid (DHA), protein ATG3, and genes *ATG7, HMGCS2* link sirtuin signaling and the PPARA network suggesting molecular mechanisms for altered hepatic gluconeogenesis from consumption of a HFr diet.

## BACKGROUND

Fructose intake in countries where people consume a Western diet has significantly increased over the past three decades, particularly through increased consumption of sweetened beverages and foods containing high-fructose corn syrup. Fructose consumption comprises a significant proportion of energy intake in the American diet, and increased consumption coincides with increased prevalence of obesity over the past three decades (1). Animal studies have shown that diets high in fructose consistently induce metabolic perturbations associated with metabolic syndrome and diabetes (1, 2). Altered metabolism in the liver has been implicated in multiple chronic metabolic diseases (3). Several studies have investigated HFr diet challenges in humans (4, 5) and nonhuman primates (NHP) (6–9). In cynomolgus monkeys (*Macaca fascicularis*), long-term exposure to high fructose (HFr) diets increased liver steatosis, with extent related to duration of fructose exposure (10), but questions remain about the initial molecular changes induced by high levels of fructose that result in long-term health complications.

The vervet monkey (*Chlorocebus aethiops sabaeus*) is a model for multiple human complex diseases including neurodegenerative disease (11), Alzheimer’s disease (12–15), diabetes, obesity and metabolism (16–18) and cardiovascular disease (19, 20) among others. Due to the high degree of genomic (21–23), physiologic and metabolic conservation between vervets and humans, results in vervets are translatable to understanding human health and disease. The ability to control environmental factors including diet and feasibility of collecting tissue biopsy samples from healthy animals, provide opportunities to investigate molecular mechanisms that are dysregulated prior to evidence of clinical disease. Studies in vervets related to metabolism have included diet interventions with variation in sources of protein, fat, and carbohydrate (18, 24, 25); However, none of these studies in humans or NHP have used global untargeted omics approaches to identify potential molecular mechanisms underlying diet-induced changes in liver metabolism. In addition, no studies to date have generated an integrated comprehensive multi-omics dataset to better understand these molecular changes (26).

The goal of this study included examination of the impact of a short-term exposure to a HFr diet in the liver, a key organ mediating carbohydrate and lipid metabolism, by integrating high-throughput omics data and investigating the benefits of data integration across multiple omics domains. The short-term HFr diet exposure has no discernible impact on body weight, insulin sensitivity, blood pressure, or triglycerides. Total plasma cholesterol and measures of liver injury were greater in animals fed the HFr diet than controls. We examined whether early molecular alterations in liver can be detected prior to development of obesity and diabetes. We compared transcriptome, proteome, and metabolome data from livers of vervets challenged with a HFr diet for six weeks with those fed a chow diet. We demonstrate that the molecular information obtained from integrated analysis of multi-omics datasets is more informative than analyses of any of the individual omics datasets. In addition, using this integrated omics approach, we identified sirtuin signaling and a peroxisome proliferator activated receptor alpha (PPARA) regulatory network as central to the hepatic short-term response to a HFr diet. Metabolites aspartic acid and DHA provide direct evidence on alterations in liver metabolism, and connect sirtuin signaling pathway and PPARA regulatory network, suggesting perturbations in these molecular mechanisms underlie altered hepatic gluconeogenesis in response to a short-term HFr diet.

## RESULTS

### Clinical and Morphometric Data Analysis

Female age-matched vervet monkeys were fed a chow diet (controls, n=5) or a HFr diet (n=5) for six weeks. Morphometric measures at the end of challenge were not different between groups. Total plasma cholesterol was increased, and measures of liver injury, alanine aminotransferase, alkaline phosphatase, and gamma-glutamyl transpeptidase were increased in animals fed the HFr diet compared to controls (Table 1).

**Table 1:**
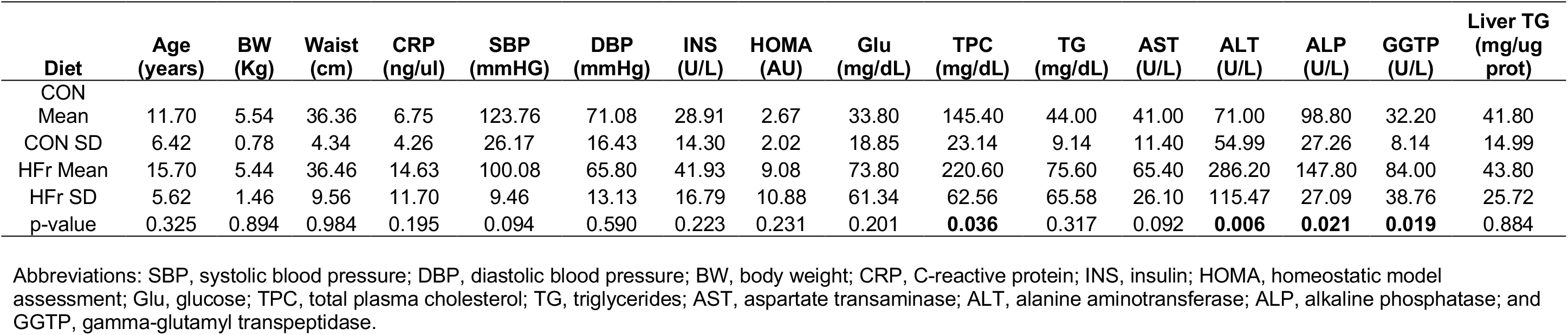
Morphometric and Clinical Measures.

### Transcriptomics Data Analysis

Comprehensive analysis of RNA expression has commonly been used to study the influence of genetic factors on phenotypic variation and is often used as a surrogate measure for functional alterations (potentially mediated by proteins or by alterations in metabolite levels). As a first step of our multi-omics characterization of liver biopsies from animals in this study, we performed RNA-Seq analyses on all samples. We identified 10,688 transcripts that passed quality filters. Of these, 467 were differentially expressed between liver samples from animals fed HFr and chow diets (unadjusted p < 0.05) (Additional file 3). Pathway enrichment analysis revealed that 51 pathways were different between HFr and chow including sirtuin signaling, remodeling of epithelial adherens junctions signaling, and necroptosis signaling (*p*-value < 0.05, Table 2, Additional files 1 and 4). Regulatory network analysis resulted in 5 networks with predicted activation states. Four networks regulated by XBP1, PPARA, MITF, and KLF15 were predicted to activate downstream targets, and one network regulated by HDAC1 was predicted to inhibit downstream targets (*p*-value < 0.05) (Table 3, Additional files 2 and 5). Regulators XBP1, PPARA, MITF, KLF15, and HDAC1 were expressed but not different between liver samples from HFr and chow-fed animals.

**Table 2:**
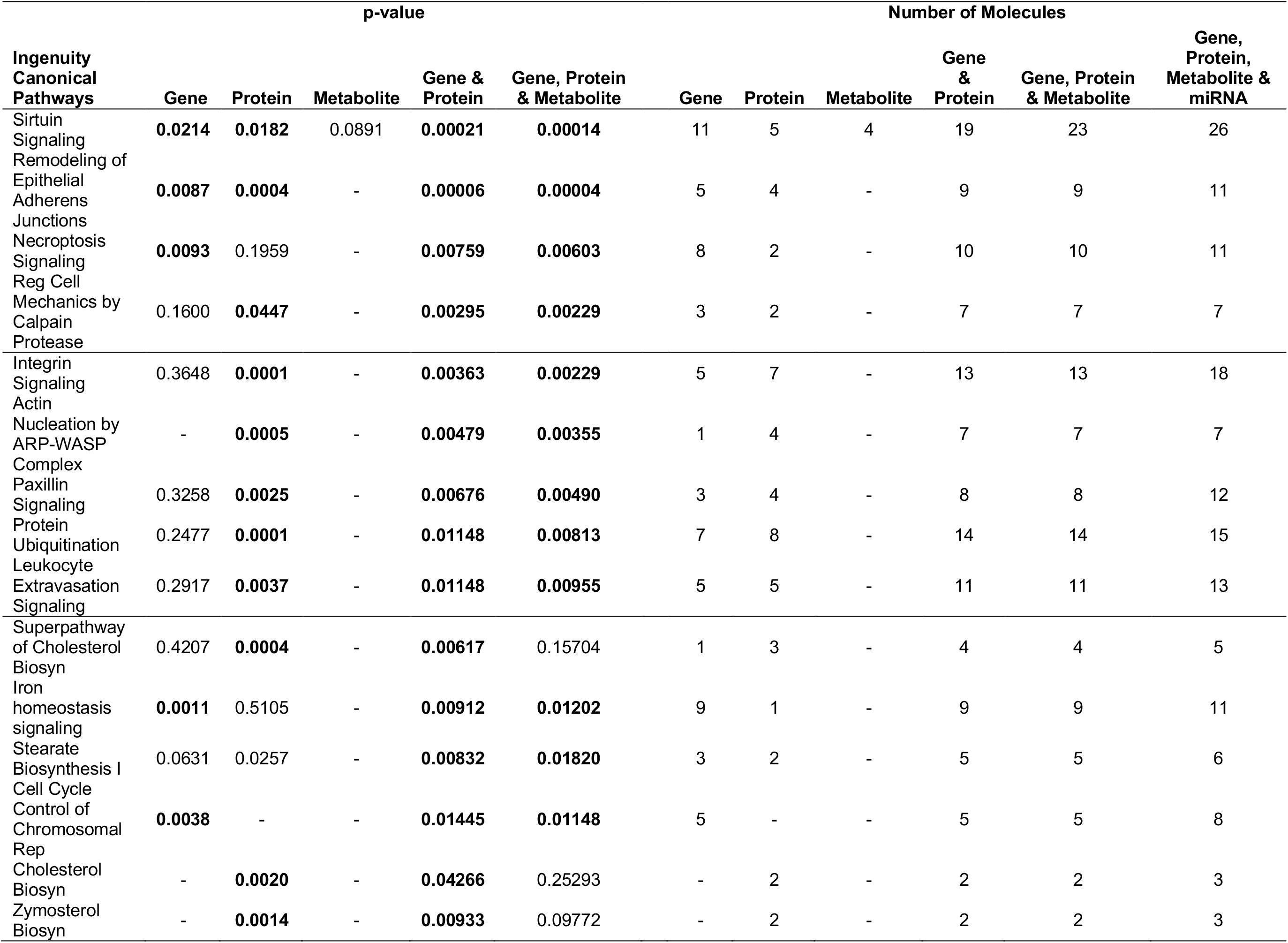
Pathways for each omic data type and integrated omics data.

| Ingenuity<br>Canonical<br>Pathways | p-value |  |  |  |  | Number of Molecules |  |  |  |  |  |
| --- | --- | --- | --- | --- | --- | --- | --- | --- | --- | --- | --- |
|  | Gene | Protein | Metabolite | Gene & Protein | Gene, Protein & Metabolite | Gene | Protein | Metabolite | Gene & Protein | Gene, Protein & Metabolite | Gene, Protein, Metabolite & miRNA |
| Sirtuin Signaling | **0.0214** | **0.0182** | 0.0891 | **0.00021** | **0.00014** | 11 | 5 | 4 | 19 | 23 | 26 |
| Remodeling of Epithelial Adherens Junctions | **0.0087** | **0.0004** | - | **0.00006** | **0.00004** | 5 | 4 | - | 9 | 9 | 11 |
| Necroptosis Signaling | **0.0093** | 0.1959 | - | **0.00759** | **0.00603** | 8 | 2 | - | 10 | 10 | 11 |
| Reg Cell Mechanics by Calpain Protease | 0.1600 | **0.0447** | - | **0.00295** | **0.00229** | 3 | 2 | - | 7 | 7 | 7 |
| Integrin Signaling | 0.3648 | **0.0001** | - | **0.00363** | **0.00229** | 5 | 7 | - | 13 | 13 | 18 |
| Actin Nucleation by ARP-WASP Complex | - | **0.0005** | - | **0.00479** | **0.00355** | 1 | 4 | - | 7 | 7 | 7 |
| Paxillin Signaling | 0.3258 | **0.0025** | - | **0.00676** | **0.00490** | 3 | 4 | - | 8 | 8 | 12 |
| Protein Ubiquitination | 0.2477 | **0.0001** | - | **0.01148** | **0.00813** | 7 | 8 | - | 14 | 14 | 15 |
| Leukocyte Extravasation Signaling | 0.2917 | **0.0037** | - | **0.01148** | **0.00955** | 5 | 5 | - | 11 | 11 | 13 |
| Superpathway of Cholesterol Biosyn | 0.4207 | **0.0004** | - | **0.00617** | 0.15704 | 1 | 3 | - | 4 | 4 | 5 |
| Iron homeostasis signaling | **0.0011** | 0.5105 | - | **0.00912** | **0.01202** | 9 | 1 | - | 9 | 9 | 11 |
| Stearate Biosynthesis I | 0.0631 | 0.0257 | - | **0.00832** | **0.01820** | 3 | 2 | - | 5 | 5 | 6 |
| Cell Cycle Control of Chromosomal Rep | **0.0038** | - | - | **0.01445** | **0.01148** | 5 | - | - | 5 | 5 | 8 |
| Cholesterol Biosyn | - | **0.0020** | - | **0.04266** | 0.25293 | - | 2 | - | 2 | 2 | 3 |
| Zymosterol Biosyn | - | **0.0014** | - | **0.00933** | 0.09772 | - | 2 | - | 2 | 2 | 3 |

**Table 3:**
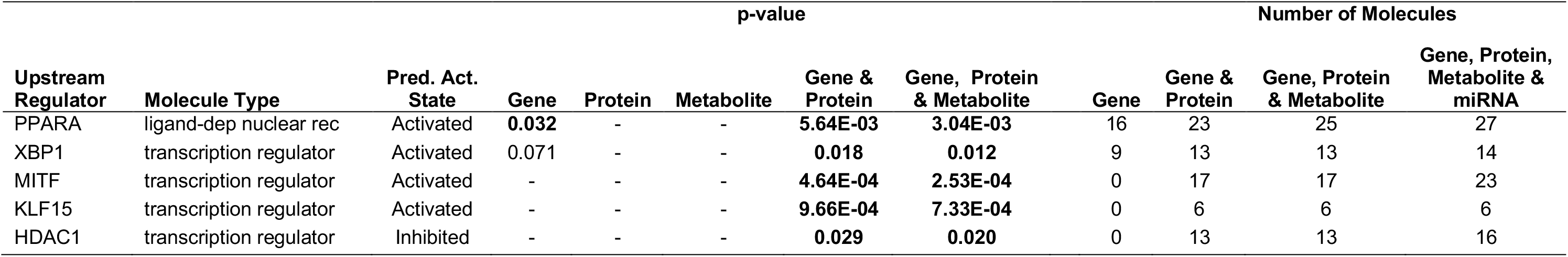
Regulatory Networks for each omic data type and integrated omics data.

### Proteomics Data Analysis

We analyzed liver-extracted proteins using standard mass spectrometry approaches as reported previously (27). Overall, we were able to identify 2858 proteins across the 10 samples. Of these, 1594 proteins were identified in at least 3 of 5 samples from either the chow- or the HFr-fed animals, and 1172 proteins were identified in samples from at least 3 animals in each group. We included further analyses the 1172 proteins plus 70 proteins that passed quality filters for all samples in one group, but were not found in any of the samples of the other group. Of the combined 1242 proteins that passed these filters, 126 proteins were quantitatively different between liver samples from HFr- and chow-fed animals (*p*-value < 0.05) (Additional file 6). Pathway enrichment analysis revealed 58 pathways altered by HFr and included pathways that were also observed from the transcriptomic data, including sirtuin signaling, and remodeling of epithelial adherens junctions signaling (*p*-value < 0.05, Table 2, Additional file 7). No regulatory networks were found with a predicted activation state (Table 3, Additional file 8). Network regulators XBP1, PPARA, MITF, KLF15, and HDAC1 were not detected in the proteomic analysis.

### Commonalities between Gene and Protein Expression

Comparison of gene and protein expression showed 320 molecules with greater expression and 263 with reduced expression that were common to both the transcriptomics and proteomics analyses in liver samples from animals fed a HFr diet compared to chow-fed animals. Comparison of statistically significant differentially expressed genes and proteins revealed only 2 shared molecules, SLCO1B1 and HTATIP2, with decreased abundance in livers from HFr-fed animals compared to chow-fed animals (Figure 1, Additional file 9).

**Figure 1:**
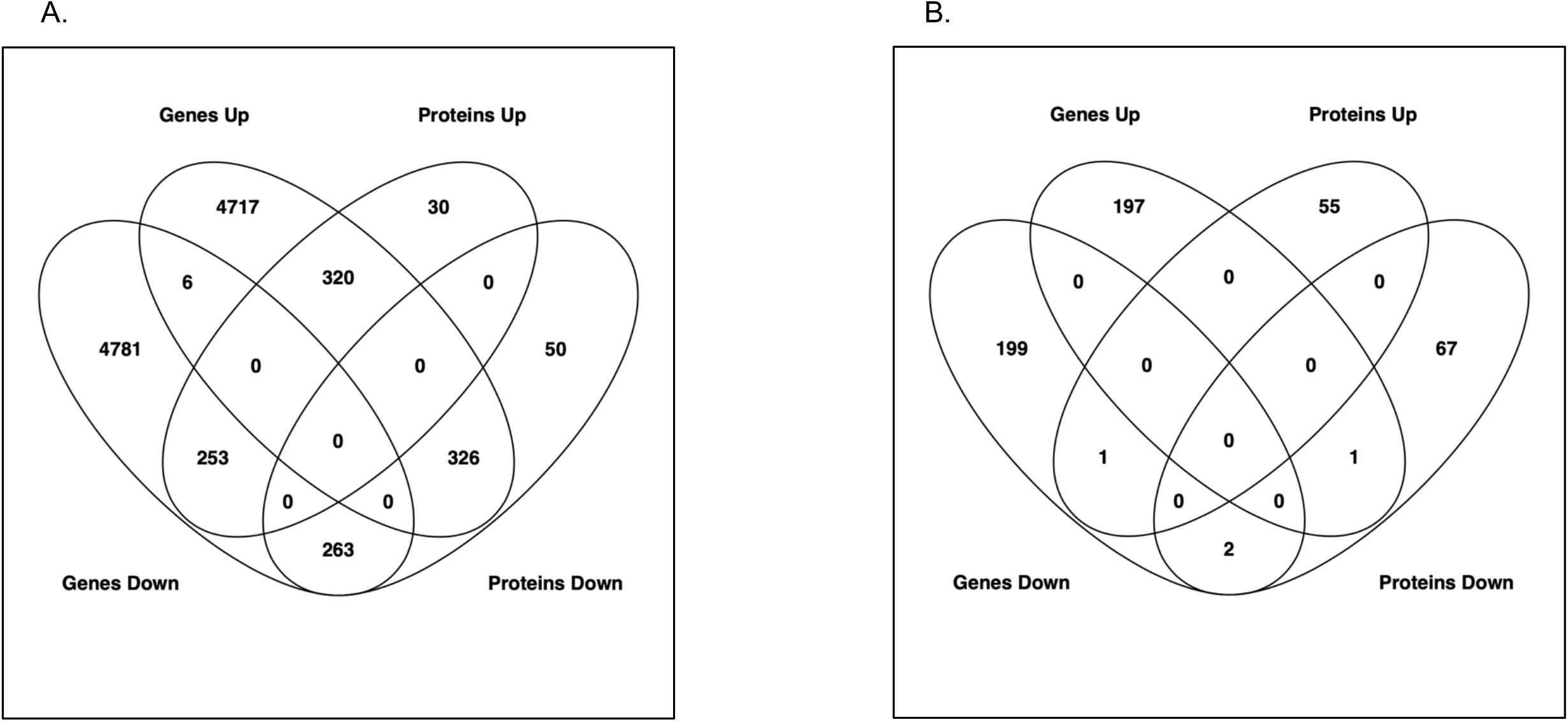
Venn diagram showing common A) expressed and B) differentially expressed genes and proteins.

### Metabolomics Data Analysis

To examine whether we could expand on the molecular changes induced in the liver by HFr exposure that we uncovered by gene-centric analyses (transcriptomics, proteomics), we performed untargeted analysis of small molecule metabolites to analyze the metabolomic changes. Overall, we quantified 471 metabolites that passed quality filters. Of these, 18 showed significantly different abundances between liver samples from HFr- and chow-fed animals (*p*-value < 0.05, Additional file 10). Pathway enrichment showed 25 pathways including aspartate biosynthesis. Sirtuin signaling was observed but not significant (*p*-value = 0.089, Table 2 and Additional file 11). All pathways identified in the enrichment analysis only contained one single metabolite per pathway, highlighting the limited annotation of metabolites in pathways and networks. No regulatory networks were found with a predicted activation state and *p*-value < 0.05 (Table 3, Additional file 12).

### Integrated Omics Analysis

Using the datasets described above, we further assessed whether combinations of omics datasets improved statistical confidence and significance in the network and pathway enrichment findings. First, we examined the combination of the gene expression and proteomics results. Integrated analysis of transcriptomic and proteomic data revealed 51 significantly enriched pathways (*p*-value < 0.05). Statistical significance of sirtuin signaling, remodeling of epithelial adherens junctions, necroptosis signaling, and regulatory cell mechanics by calpain protease increased, and the number of molecules identified in each network increased with dataset integration. Interestingly, for sirtuin signaling, the number of genes and proteins was greater than the sum of genes and proteins from individual omic pathway analysis; this is due to our requirement for direct connections with addition of protein data to gene data connecting additional genes in the pathway. Significance of some pathways decreased, such as stearate biosynthesis, cell cycle control of chromosomal replication, and cholesterol biosynthesis (Table 2, Additional files 1 and 13). Integrated analysis showed 4 activated networks with predicted regulators PPARA, XBP1, MITF, and KLF15, and one inhibited network with predicted regulator HDAC1. Statistical significance increased and the number of molecules in the networks increased for the PPARA and XBP1 networks when compared to the analysis of the transcriptomic data alone (Table 3, Additional files 2 and 14).

Integration of the transcriptomics and proteomics data with metabolomics findings further enhanced the pathway enrichment and network analyses, and resulted in the identification of 43 significantly enriched pathways. The significance of several pathways, and the number of molecules identified in each pathway, increased even more compared to the gene-protein integrated pathways, including again sirtuin signaling, remodeling of epithelial adherens junctions, necroptosis signaling, and regulatory cell mechanics by calpain protease. Sirtuin signaling had the greatest significance and the greatest number of identified molecules with genes, proteins and metabolites. In addition, significance of other pathways such as cell cycle control of chromosomal replication, and cholesterol biosynthesis further decreased again when compared to the gene-protein integrated networks (Table 2, Additional files 1 and 15). Integrated network analysis was similar to pathway analysis with increased significance and molecule number compared to the gene-protein integrated networks, with the PPARA regulatory network (that included gene transcripts, proteins and metabolites) being the most significant (Table 3, Additional files 2 and 16). Of note, the protein FASN directly links regulatory networks PPARA, XBP1 and KLF15. In addition, overlapping molecules in networks link regulators PPARA and KLF15 with sirtuin signaling, including the protein ATG3, gene transcripts ATG7, HMGCS2, and metabolites DHA and L-aspartic acid (Figure 2).

**Figure 2:**
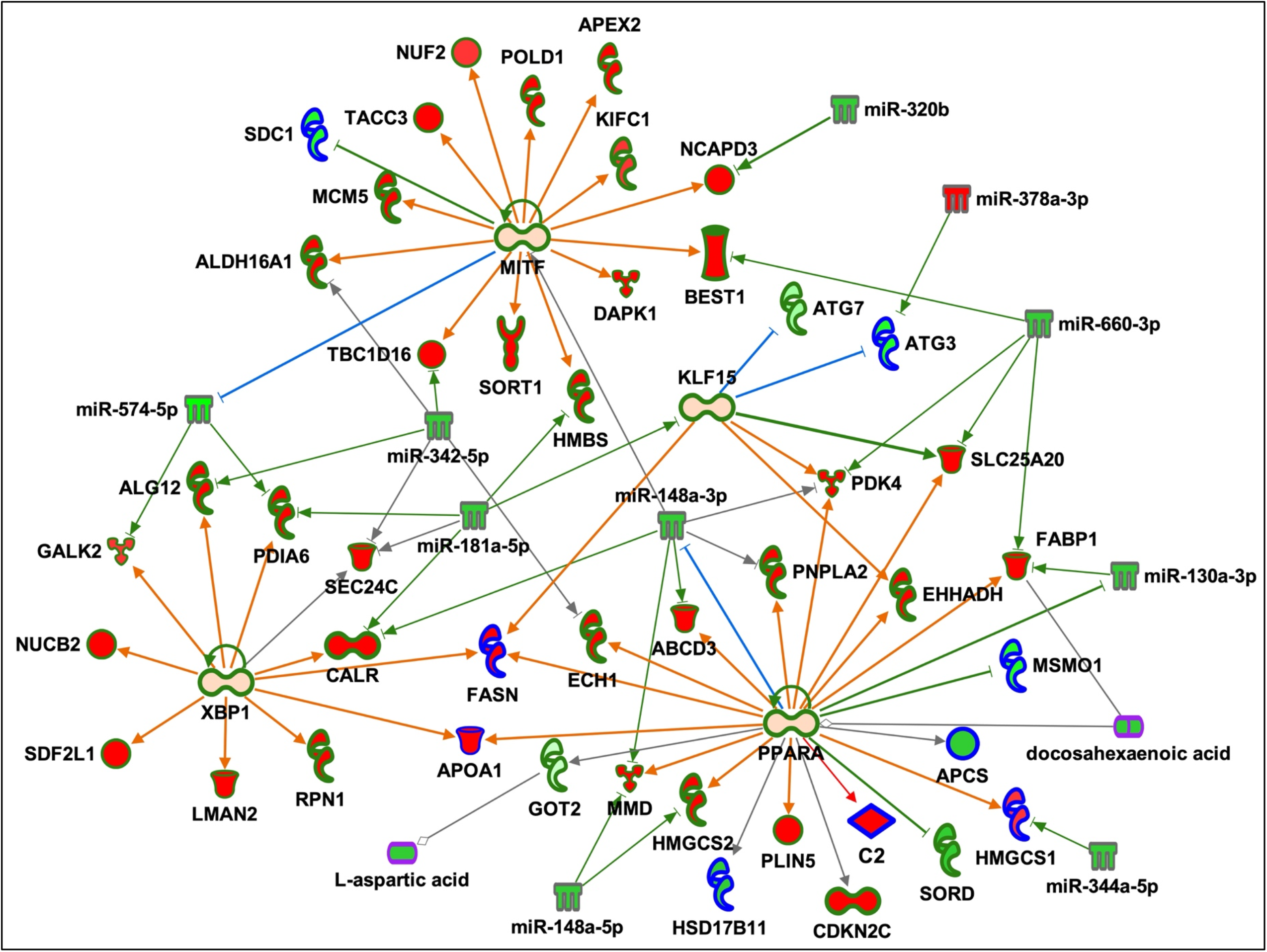
Regulatory network up-regulated in HFr livers compared with chow. Red fill indicates increased abundance, green fill decreased abundance, light orange fill indicates predicted activation, green outline genes, blue outline proteins, gray outline miRNAs, purple outline metabolites, green lines indicate inhibition and red lines activation.

### Integration of miRNA Data

In an effort to explore putative regulatory mechanisms underlying the pathway and network enrichment we describe above, we integrated analysis data from small RNA-Seq (which characterizes miRNAs) with the multi-omics datasets described above. In our analysis, we identified 576 known miRNAs that passed quality filters. Of these, 22 were differentially expressed between liver samples from HFr- and chow-fed animals (*p*-value < 0.05, Additional file 17). Detailed miRNA – gene/protein pairing provided a list of 793 inverse pairs that included 17 differentially expressed miRNAs and 758 differentially abundant genes or proteins (Additional file 18). Integration of miRNAs with pathways increased the number of molecules in remodeling of epithelial adherens junctions and necroptosis signaling, and the number of molecules increased for regulatory networks PPARA, XBP1, MITF and HDAC1 (Table 3, Additional file 2). In addition, these regulatory networks were interconnected by miRNAs that target genes and proteins in multiple networks: miR-148-3p for PPARA, MITF, KLF15, and XBP1 network genes and proteins, miR-181a-5p for MITF, KLF15, and XBP2 network genes and proteins, miR 342-5p for MITF, XBP1 and PPARA network genes and proteins, and miR-574-5p for XBP1 and MITF network genes and proteins (Figure 2). This integration suggests potential regulatory roles for these miRNAs in coordinating the molecular changes induced in the liver after exposure to a HFr diet, and emphasizes the complexity of miRNA interactions that may affect both transcript and protein levels.

### Genes and Proteins in Multi-Omic Networks with Associations to NASH- and NAFLD-Related Traits

To examine the potential shared pathophysiological mechanisms induced by short term HFr diet exposure with long-term liver health outcomes associated with HFr, we compared GWAS catalog variants and genes associated with nonalcoholic steatohepatitis (NASH)- and nonalcoholic fatty liver disease (NAFLD)-related traits, including BMI, lipoproteins, obesity, diabetes, insulin resistance, with the differentially expressed genes and proteins identified in our analysis of liver samples. The alignment of the datasets revealed 53 genes and proteins with one or more intergenic single nucleotide polymorphism (SNP) associated with one or more NASH/NAFLD related trait(s) (Additional file 19). When we restricted the analysis only to genes and proteins in significantly enriched multi-omic pathways and networks, we identified 13 genes with GWAS SNPs, including FABP1 (associated with NAFLD) in PPARA and HDAC1 networks; GOT2 (associated with triglycerides and aspartate aminotransferase) in the sirtuin signaling pathway; and ATG7 (associated with fat body mass) in the sirtuin signaling pathway and KLF15 network (Table 4).

**Table 4:** Pathway and Network Genes and Proteins with GWAS SNPs.

| **Gene**<br>**Symbol** | **Pathway or Network** | **Trait** |
| --- | --- | --- |
| APOA1 | HDAC1 Network<br>PPARA Network<br>XBP1 Network | Very low-density lipoprotein cholesterol |
| ATG7 | KLF15 Network<br>Sirtuin Signaling Pathway | Fat body mass |
| CLIP1 | Remodeling of Epithelial Adherens Junctions | Body mass index |
| FABP1 | HDAC1 Network<br>PPARA Network | Non-alcoholic fatty liver disease<br>Hepatic fibrosis |
| GOT2 | Sirtuin Signaling Pathway | Triglycerides<br>Aspartate aminotransferase |
| MET | MITF Network<br>Remodeling of Epithelial Adherens Junctions | Triglycerides |
| MITF | MITF Network | Low-density lipoprotein cholesterol<br>Triglycerides |
| PNPLA2 | PPARA Network | Body fat distribution |
| PPARA | PPARA Network | Type II Diabetes<br>Total cholesterol<br>Low-density lipoprotein cholesterol<br>Triglycerides |
| RAC1 | Actin Nucleation by ARP-WASP Complex<br>Integrin Signaling<br>Leukocyte Extravasation Signaling<br>Paxillin Signaling | Low-density lipoprotein cholesterol |
| RAP1GAP | Leukocyte Extravasation Signaling | Alkaline phosphatase |
| SORT1 | MITF Network | Type II Diabetes<br>Coronary artery disease<br>LDL cholesterol change |
| TNFRSF11B | Necroptosis Signaling Pathway | Alkaline phosphatase |

## DISCUSSION

The liver is central to metabolic regulation, and dysregulation of liver metabolism directly impacts gluconeogenesis and lipogenesis. Exposure to a HFr diet is known to increase the risk of dyslipidemia, insulin resistance, lipogenesis (28), levels of hepatic oxidative stress markers, and induce NASH and NAFLD (6). Unlike glucose, fructose is absorbed in the intestine independently of energy or sodium exchange. When consumed in high amounts, fructose is transported to the liver via hepatic portal circulation and is preferentially converted to lipids. Fructose forms the building blocks of triglycerides (29), and triglycerides produced in the liver mostly are packaged into atherogenic very low-density lipoprotein particles (30). Fructose in the liver can also serve as substrate for the gluconeogenesis pathway and increase circulating glucose levels (31), which, together with the increased triglyceride levels, decreases overall glycemic control. The specific contribution of hepatic steatosis to whole body insulin sensitivity and dyslipidemia (32–35) is particularly significant for individuals diagnosed with the metabolic syndrome. However, the underlying molecular networks that are dysregulated by a HFr diet and precede insulin resistance, NASH and NAFLD have not yet been identified, and the initial molecular abnormalities initiated by the exposure to fructose remain to be identified (6).

NHPs have been shown to be valuable models of diet-induced metabolic dysregulation due to extensive similarities with human metabolism (7). The ability to carefully control diet exposure, and the physiological similarity to humans make NHP an ideal model to examine molecular tissue and organ changes in response to short- and long-term dietary challenges. We used a cohort of vervet monkeys (*Clorocebus aethiops sabeus*) fed an acute HFr diet (n=5) or chow diet (n=5) for 6 weeks. Previous analyses showed changes in liver enzymes, total plasma cholesterol, and liver histology indicative of liver injury with periportal and inflammatory lesions in the HFr group (6), but no other clinically discernable abnormalities in body mass, or circulating glucose levels. In this study, we used global untargeted transcriptomics, proteomics, and metabolomics of liver biopsy samples to identify the acute early hepatic molecular and cellular response to a HFr diet, prior to onset of fat accumulation or systemic pathophysiological changes, to identify dysregulated molecular networks that potentially drive fat accumulation, and may be the initiating steps for subsequent long-term liver dysregulation. Pathway and network analyses were performed on individual datasets and integrated multi-omics datasets to determine whether there was a gain in our understanding of the molecular impact of a HFr diet with a combined approach compared to use of single or double omics datasets. Our analytical approach included prioritization of molecules by using pathway and network enrichment statistics, with the stringent requirement of direct connections among molecules, to improve statistical rigor for this study with small sample sizes (a common limitation of NHP studies).

We chose to use IPA to assess integrated omics effectiveness since it has tools for canonical pathway enrichment, and the underlying knowledgebase provides a means for regulatory network analysis at high resolution using transcripts, proteins, and metabolites, which is not yet feasible with other publicly available tools such as DAVID Bioinformatic Resources (36). Our findings confirm previous papers indicating the need for better tools to perform integrated omic analyses (26). In addition, it will be important to test strengths and limitations of multi-omics data integration with other tools when available.

In analyzing individual omics datasets, we identified a large number of statistically significant pathways for each data type, which is often the case for these types of data, making it a challenge to prioritize networks and distinguish likely true associations from spurious results. Integration of hepatic transcriptomic and proteomic data increased the significance of a number of pathways and networks, while decreasing the significance of other pathways, suggesting that truly associated pathways can be distinguished better with this approach. Interestingly, comparison of differentially expressed genes and proteins showed very little overlap: potentially due to the low correlation usually observed in expressed protein and transcript abundances. Most studies investigating proteome and transcriptome in the same model have noted this (e.g. (37)). However, integration of these datasets provided additional molecules with direct connections within a pathway or network, increasing the overall number of molecules, increasing the confidence in pathway or network prediction, and providing additional information about molecular functions. For some pathways and networks, additional differentially abundant molecules were added from the second omics dataset, creating new connections not evident in either of the individual omics datasets. Of note, proteins are often identified as molecules connecting separate regulatory networks and steps within signaling pathways, e.g. ATG3 in sirtuin signaling and FASN for the XBP1, PPARA and KLF15 networks.

Integration of transcriptomic and proteomic data increased the significance of the sirtuin signaling pathway, and revealed direct connections between sirtuin signaling and the four activated networks with predicted regulators PPARA, XBP1, MITF and KLF15. It is important to note that all of these genes were detected but not differentially expressed, but the encoded proteins were not detected. These results do not contradict the role of these proteins as central regulators since activity of all four depend on post-translational modifications (38–43).

Integration of metabolomic data with transcriptomic and proteomic datasets further improved significance of some pathways, with sirtuin signaling increasing in rank and statistics from being 7^th^ for transcriptomics and 39^th^ for proteomics, to becoming 2^nd^ for transcriptomics and proteomics, and 2^nd^ overall with integration of all 3 datatypes. This pathway included the most molecules, including 4 metabolites. Other pathways decreased in significance and rank compared with the analysis of individual omics datasets. Addition of metabolites also provided more direct connections among regulatory networks, and connected the sirtuin signaling pathway with the PPARA network. Metabolites aspartic acid and DHA also indicated end-of-pathway directionality for the sirtuin signaling pathway and the PPARA network.

Finally, integration of miRNA data showed 19 of 22 differentially expressed miRNAs targeted genes and/or proteins in the four activated networks and sirtuin signaling pathway with inverse expression profiles. Our miRNA findings suggest that the initial hepatic response to short-term exposure to a HFr diet is at least in part epigenetically regulated. Taken together, these results demonstrate that integration of transcriptomic, small transcriptomic, proteomic, and metabolomic data reveals pathways and networks central the HFr diet response in the liver, not seen by analysis of only one or two of these omic datasets.

Our results from these unique NHP biopsy samples reveal interesting novel molecular mechanisms regulating the initial hepatic response to HFr diet exposure in these animals. The sirtuin signaling pathway and networks regulated by PPARA, XBP1, MITF and KLF15 appear to be central to the HFr diet response. Both sirtuin signaling (44, 45) and PPARA (46) play important roles in the pathophysiology of NAFLD. For the sirtuin gene family, the majority of studies have focused on the role of SIRT1 in regulating both lipid and carbohydrate metabolism (47–49). Interestingly, in our study, SIRT2 rather than SIRT1 was central to the initial hepatic response to a HFr diet. A recent study in male mice showed that SIRT2 functions as a negative regulator of NAFLD development and progression, with increased expression being protective when animals were fed a high-fat diet (50). Our study in female NHPs showed higher SIRT2 expression in the HFr group compared with chow-fed animals, and lower expression of GOT2 and decreased abundance of aspartic acid (51), which is regulated by GOT2 (52, 53). In mice, quantification of GOT2 protein expression by immunohistochemistry shows decreased abundance with NAFLD (54), supporting our preliminary findings. GOT2 and aspartic acid are at the end of the sirtuin pathway and indicative of altered gluconeogenesis and pathologies associated with NAFLD.

While the overall pathways identified in our analysis are supported by published evidence in other model organisms and related pathophysiologies, we also raise additional questions about previously under- or unappreciated regulatory networks. Our analysis suggests that the HFr diet exposure led to activation of the PPARA network, and downstream molecules GOT2 and aspartic acid showed decreased abundance. Studies of PPARA liver expression in mice with steatosis in response to a high-fat diet show sex-differences: PPARA expression is increased in male rats, and FASN, which is directly downstream of PPARA, is also increased. However, in female rats, FASN is increased but PPARA is not (55), suggesting that hepatic PPARA activation/inhibition of FASN may be sex-specific, and the potentially divergent expression patterns in our female NHP in response to the HFr diet may be specific to female animals.

As another example, our detailed multi-omics analysis also suggested that DHA, an omega-3 polyunsaturated fatty acid with anti-inflammatory functions (56), was lower in livers from animals fed a HFr diet than in livers from chow-fed animals. While no studies have reported changes in DHA in response to fructose, human studies examining dietary supplementation with DHA have suggested the beneficial effects of the increased level of DHA may include decreased incidence of NAFLD (57). DHA is known to bind and activate PPARA (58) which may influence sirtuin signaling and the integrated regulatory network we discovered in our analysis. The decreased abundance of DHA, but with predicted activation of PPARA and activation of all but GOT2 downstream of PPARA, like aspartic acid, suggests differences between rodents and primates or sex-differences in these signaling networks, and may point to other mechanisms (apart from DHA) by which PPARA expression may be increased by HFr.

GWAS of genes and proteins in sirtuin signaling and the four activated networks we identified show a single gene, FABP1, that has been reported to be associated with alanine aminotransferase levels, a marker of liver disease (59). Twelve additional genes were associated with lipoprotein-, insulin-, and BMI-related traits. Identification of SIRT2 and an integrated network of regulatory genes and proteins with altered abundance in livers from animals exposed to a HFr diet that are upstream of GOT2 and aspartic acid suggest that we have identified novel molecules and regulatory mechanisms that influence and potentially govern the initial hepatic response to short-term HFr diet exposure. Additional studies are required to validate our findings, and to explore potential targets by which these networks can be modulated to blunt the effects of fructose consumption on overall liver metabolism and function, preventing subsequent health complications known to occur with high intake levels.

## CONCLUSIONS

We have demonstrated that integration of multiple omics datasets significantly improves prioritization of pathways and networks that influence hepatic response to a short-term HFr diet. Using this integrated approach, we identified sirtuin signaling and a large, integrated regulatory network, with molecules overlapping sirtuin signaling as a potential key modulator and regulator of hepatic metabolism in response to a HFr diet.

## MATERIALS AND METHODS

### Animals and Experimental Design

All experimental procedures involving vervet monkeys (*Chlorocebus sabaeus*) were approved and complied with the guidelines of the Institutional Animal Care and Use Committee of Wake Forest University Health Sciences, which is an AALAC accredited facility. Procedures were performed by a veterinarian (KK), including liver biopsy as previously described (27). Animals were provided non-steroidal anti-inflammatory and opioid analgesics during recovery as needed. Liver tissue was flash frozen in liquid nitrogen and stored at −80 C until analysis. Animal housing, handling, diet compositions (chow and HFr) and caloric details are as described elsewhere (6). Prior to the study, all animals were maintained on chow diet. For this study, 10 female vervet monkeys were fed with either chow (n=5) or HFr (n=5) diets for 6 weeks. Previous studies have shown sex-specific metabolic responses to a HFr diet (7); for this reason, all animals in the study were female.

### Clinical Measures

Serum-based clinical measures, including total protein, albumin, globulin, albumin/globulin ratio, AST, ALT, ALK phosphatase, GGTP, total bilirubin, urea nitrogen, creatinine, BUN/creatinine ratio, phosphorus, glucose, calcium, magnesium, sodium, potassium, Na/K ratio, chloride, cholesterol, triglycerides, amylase, lipase, CPK, and hematological parameters including WBC, RBC, hemoglobin, hematocrit, MCV, MCH, MCHC, blood parasites, platelet count, platelet, EST, neutrophils, bands, lymphocytes, monocytes, eosinophils and basophil data were obtained from ANTECH Diagnostics (800-872-1001, NC, USA).

### Transcriptomics: RNA Seq

#### RNA Extractions and Sequencing

Total RNA was extracted from vervet monkey livers using the Zymo Direct-zol™ kit (Zymo Research) and each sample was subsequently quantified by Qubit assay (Thermo Fisher). RNA-Seq libraries were prepared from 500 ng of total RNA according to the Illumina TruSeq stranded mRNA protocol (Illumina), which specifically retains polyadenylated mRNAs by the oligo dT coated magnetic beads. Sequencing library concentrations were quantified using the KAPA library quantification kit (Kapa Biosystems). Clusters were generated by cBot (Illumina), and 2 × 100 base paired-end sequencing libraries were sequenced using the Illumina HiSeq 2500 with v3 sequencing reagents (Illumina).

#### Data Analysis

Raw sequences were de-multiplexed using the Illumina pipeline CASAVA v1.8. The FastQC and FASTX toolkit were used for QC. Sequence reads with Phred scores ≥ Q30 were retained. Reads aligned against the vervet reference genome (ChlSab1.1) were annotated using the Ensembl release 93 gene model. Abundance analysis was performed using our established RNA-Seq workflow in Partek Flow, which allowed calculation of transcript-level expression of a gene’s isoforms for alternative spliced transcripts (60, 61). Transcript abundances were quantified in Flow (Partek) using an expectation-maximization algorithm similar to the reported (62) which quantifies isoform expression levels across the whole genome at the same time and normalizes by transcript length to account for the transcript fragmentation step in RNA-Seq. Transcripts without read counts across all samples were filtered out, and then normalized by the trimmed mean of M values method [Robinson MD and Oshlack A. Genome Biol. 11:R25, 2010] Differentially expressed genes were identified by Analysis of Variance (ANOVA; unadjusted p < 0.05). Gene expression data were deposited in the National Center for Biotechnology Information’s Gene Expression Omnibus (GEO; http://www.ncbi.nlm.nih.gov/geo/) - GEO Series accession number GSE176576.

### Transcriptomics: small RNA Seq

#### Sequencing

RNA extracted for RNA Seq was also used for small RNA Seq. Small RNA Seq methods are described in (63). Briefly, small RNA sequencing libraries were prepared using the Illumina TruSeq Small RNA Sample Prep Kit and were pooled after cDNA synthesis. cDNA libraries were clustered using an Illumina Cluster Station and sequenced with an Illumina GAIIx sequencer. Raw sequence reads were obtained using Illumina’s Pipeline v1.5. Extracted sequence reads were normalized, annotated and abundance determined using mirDeep2 (64).

#### Data Analysis

Transcripts without read counts across all samples were filtered out, and then normalized by the trimmed mean of M values method. Differentially expressed genes identified by Analysis of Variance (ANOVA; unadjusted p < 0.05). Gene expression data were deposited in the National Center for Biotechnology Information’s Gene Expression Omnibus (GEO; http://www.ncbi.nlm.nih.gov/geo/) - GEO accession number GSE178269.

### Proteomics

Proteomics data were generated by liquid chromatography-coupled tandem mass spectrometry using a Thermo Scientific Orbitrap Elite mass spectrometer. Details of sample preparation, mass spectral analysis, and data analysis using a proteogenomics approach in Morpheus were described previously (27).

#### Data Analysis

For each animal, peptide spectrum intensities reported in Morpheus were summed across occurrences (i.e. across multiple transcript matches) based on Gene IDs. Proteins identified and quantified in at least 3 animals per group (HFr and chow) retained for downstream analysis. Additionally, proteins that were quantified in all samples of one group but not in any of the samples of other group were also retained for subsequent analyses. Intensity values were log transformed, and missing data (at most 2 animals per group) were imputed using the NAguideR tool with the impseq approach (sequential imputation) separately for the two experimental groups (HFr or chow).

#### Comparison of gene and protein abundance

Gene lists (Additional file 3) and protein lists (Additional file 4) were uploaded into Venny and Venn diagrams were generated showing commonly expressed and differentially expressed genes and proteins (65). Ratios of HFr to chow were used to determine directionality.

### Metabolomics

#### GC-TOFMS Analysis

Liver metabolites were analyzed with chemical derivatization following previously published protocols (66, 67). Extracted samples were spiked with two internal standard solutions (10 μL of L-2-chlorophenylalanine in water, 0.3 mg/mL; 10 μL of heptadecanoic acid in methanol, 1 mg/mL), mixed, and extracted with 300 μL of methanol/chloroform (3:1). After centrifugation at 12 000*g* for 10 min, an aliquot of the 300-μL supernatant was transferred to a glass sampling vial to vacuum-dry at room temperature. The residue was derivatized using a two-step procedure. First, 80 μL of methoxyamine (15 mg/mL in pyridine) was added to the vial and kept at 30 °C for 90 min, followed by 80 μL of BSTFA (1% TMCS) at 70 °C for 60 min.

Each 1-μL aliquot of the derivatized solution was injected in splitless mode into an Agilent 6890N gas chromatograph coupled with a Pegasus HT time-of-flight mass spectrometer (Leco Corporation, St. Joseph, MI). The CRC and control samples were run in the order of “control-CRC-control”, alternately, to minimize systematic analytical deviations. Separation was achieved on a DB-5ms capillary column (30 m × 250 μm i.d., 0.25-μm film thickness; (5%-phenyl)-methylpolysiloxane bonded and cross-linked; Agilent J&W Scientific, Folsom, CA), with helium as the carrier gas at a constant flow rate of 1.0 mL/min. The temperature of injection, transfer interface, and ion source was set to 270, 260, and 200 °C, respectively. The GC temperature programming was set to 2 min isothermal heating at 80 °C, followed by 10 °C/min oven temperature ramps to 180 °C, 5 °C/min to 240 °C, and 25 °C/min to 290 °C, and a final 9 min maintenance at 290 °C. Electron impact ionization (70 eV) at full scan mode (*m/z* 30-600) was used, with an acquisition rate of 20 spectra/s in the TOFMS setting.

#### GC-TOFMS Data Analysis

The acquired MS files from GC-TOFMS analysis were exported in NetCDF format by ChromaTOF software (v3.30, Leco Co., CA). CDF files were extracted using custom scripts (revised Matlab toolbox hierarchical multivariate curve resolution (H-MCR), developed (68, 69) in the MATLAB 7.0 (The MathWorks, Inc.) for data pretreatment procedures such as baseline correction, denoising, smoothing, alignment, time-window splitting, and multivariate curve resolution (based on multivariate curve resolution algorithm) (68). The resulting data set includes sample information, peak retention time and peak intensities. Compound identification was performed by comparing the mass fragments with National Institute of Standards and Technology (NIST) 05 Standard mass spectral databases in NIST MS search 2.0 (NIST, Gaithersburg, MD) software with a similarity of more than 70% and finally verified by available reference compounds.

#### 2D GC-ToF-MS Analysis

Gas chromatography-mass spectrometry was performed as described (70). Metabolite extracts were dried under vacuum in cold, and were then sequentially derivatized with methoxyamine hydrochloride (MeOX) and *N*-methyl-*N*-trimethylsilyl-trifluoroacetamide (MSTFA) (70). One microliter of the derivatized sample was injected in splitless mode using an autosampler (VCTS, Gerstel™, Linthicum, MD, USA) into a GC-MS system consisting of an Agilent^©^ 7890 B gas chromatograph (Agilent Technologies, Palo Alto, CA, USA) with Pegasus ® 4D ToF-MS instrument (LECO Corp., San Jose, CA, USA) equipped with an electron impact (EI) ionization source. Injection of the sample was performed at 250 °C with helium as a carrier gas and flow set to 2 mL min^-1^. GC was performed using a primary Rxi®-5Sil MS capillary column (Cat. No. 13623-6850, Restek, Bellefonte, PA, USA) (30 m × 0.25 mm × 0.25 μm) and a secondary Rtx®-17Sil capillary column (Cat. No. 40201-6850, Restek, Bellefonte, PA, USA). The temperature program started isothermal at 70 °C for 1 min followed by a 6 °C min^-1^ ramp to 310 °C and a final 11 min hold at 310 °C. The system was then temperature-equilibrated at 70 °C for 5 min before the next injection. Mass spectra were collected at 20 scans/s with a range of *m/z* 40-600. The transfer line and the ion source temperatures were set to 280 °C. QC standards were injected at scheduled intervals for tentative identification and monitoring shifts in retention indices (RI).

#### 2D GC-ToF-MS Data Analysis

The GC-MS data were pre-processed, cleaned, aligned, and processed using ChromaToF version 4.50.8.0 (LECO Corp., Michigan, USA) following settings from (71). Briefly described settings viz. S/N: 5; peak width: 0.15, base line offset: 1; m/z range: 50-800. The aligned data were also deconvoluted using Automated Mass Spectral Deconvolution and Identification System (AMDIS, NIST, USA) interface to match against the freely available MSRI spectral libraries of the Golm Metabolome Database available from Max-Planck-Institute for Plant Physiology, Golm, Germany (http://csbdb.mpimp-golm.mpg.de/csbdb/gmd/gmd.html) by matching the mass spectra and RI (72). Metabolites were identified by comparing fragmentation patterns available in both the Golm database as well as NIST Mass Spectral Reference Library (NIST11/2011; National Institute of Standards and Technology, USA) library. Peak finding and quantification of selective ion traces were accomplished using AMDIS software. Base peak areas of the mass fragments (*m/z*) were normalized using median normalization and log_2_ transformation. Peak areas were normalized by dividing each peak area value by the area of the internal standard for a specific sample, and were further median normalized.

#### Liquid Chromatography-Time of Flight Mass Spectrometry (LC-TOFMS)

Plasma samples were processed as reported before (73). A volume of 100 μL supernatant was mixed with 400 μL of a mixture of methanol and acetonitrile (5:3). Liver tissue homogenate was added to 500 μL of a chloroform, methanol, and water mixture (1:2:1, v/v/v). These samples were then mixed and centrifuged at 13,000 rpm for 10 min at 4°C. A 150 μL aliquot of supernatant was transferred to a sampling vial. The deposit was re-homogenized with 500 μL methanol followed by a second centrifugation. Another 150 μL supernatant was added to the same vial for drying and then reconstituted in 500 μL of ACN: H2O (6:4, v/v) before separation.

An Agilent HPLC 1200 system equipped with a binary solvent delivery manager and a sample manager (Agilent Corporation, Santa Clara, CA, USA) was used with chromatographic separations performed on a 4.6 × 150 mm 5 μm Agilent ZORBAX Eclipse XDB-C18 chromatography column. The LC elution conditions are optimized as follows: isocratic at 1% B (0–0.5 min), linear gradient from 1% to 20% B (0.5–9.0 min), 20–75% B (9.0–15.0 min), 75–100% B (15.0–18.0 min), isocratic at 100% B (18–19.5 min); linear gradient from 100% to 1% B (19.5–20.0 min) and isocratic at 1% B (20.0–25.0 min). For positive ion mode (ESI+) where A = water with 0.1% formic acid and B = acetonitrile with 0.1% formic acid, while A = water and B = acetonitrile for negative ion mode (ESI-). The column was maintained at 30 °C as a 5 μL aliquot of sample is injected. Mass spectrometry is performed using an Agilent model 6220 MSD TOF mass spectrometer equipped with a dual sprayer electrospray ionization source (Agilent Corporation, Santa Clara, CA, USA). The TOF mass spectrometer was operated with the following optimized conditions: (1) ES+ mode, capillary voltage 3500 V, nebulizer 45 psig, drying gas temperature 325 °C, drying gas flow 11 L/min, and (2) ES− mode, similar conditions as ES+ mode except the capillary voltage was adjusted to 3000 V. During metabolite profiling experiments, both plot and centroid data are acquired for each sample from 50 to 1,000 Da over a 25 min analysis time. Data generated from LC-TOFMS were centroided, deisotoped, and converted to mzData xml files using the MassHunter Qualitative Analysis Program (vB.03.01) (Agilent). Following conversion, xml files are analyzed using the open source XCMS package (v1.16.3) (http://metlin.scripps.edu), which runs in the statistical package R (v.2.9.2) (http://www.r-project.org), to pick, align, and quantify features (chromatographic events corresponding to specific m/z values and elution times). The software is used with default settings as described (http://metlin.scripps.edu) except for xset (bw = 5) and rector (plottype = “m”, family = “s”). The created .tsv file is opened using Excel software and saved as .xls file. Compound identification was performed by comparing the accurate mass and retention time with reference standards available in our laboratory, or comparing the accurate mass with online database such as the Human Metabolome Database (HMDB). Metabolomic LC/GC-TOFMS data was analyzed using principle component analysis (PCA) and OPLS analysis between groups. The differential metabolites were selected when they meet the requirements of variable importance in the projection (VIP) >1 in OPLS model and p < 0.05 from student *t*-test. The corresponding fold change shows how these selected differential metabolites varied from control. Final data analysis between control HFr-diet groups for each metabolite was conducted using independent *t*-test analysis with a *p* < 0.05 significance threshold.

### Pathway and Network Analyses

For individual omic datasets, all quality molecules for the dataset were uploaded to Ingenuity Pathway Analysis (IPA; QIAGEN). Gene symbols were used for genes and proteins, which are conserved between human and vervet. Pathway and network enrichment analyses used differentially abundant molecules and the IPA Knowledge Base, and requiring direct connections based on experimental evidence among differentially abundant molecules. Right-tailed Fisher’s exact test was used to calculate enrichment of differentially expressed genes in pathways, p< 0.01(61). Regulatory network prediction required previous experimental validation of direct connections in liver or liver cells.

### Integrated Omic Analyses

Multi-omic data analysis combined the total gene, protein, and/or metabolite lists for all molecules that passed quality filters as appropriate for the data type. Lists included molecule ID, direction of change, fold change, and p-value. Pathway and network enrichment used the same parameters and statistical tests as for individual omic datasets, requiring experimentally validated direct connections for differentially abundant molecules.

### miRNA – Gene/Protein pairing

Current pathway and network enrichment tools in IPA do not provide the means to filter direct connections based on inverse abundance between a miRNA and its target. In order to integrate our miRNA data, we performed miRNA – gene pairing in IPA for our miRNA, gene and protein datasets, requiring opposite expression for experimentally validated or highly predicted interactions (e.g., HFr miRNA up-regulated and HFr gene down-regulated compared with chow). Using the gene and protein IDs in this list, we merged it with the list of genes and proteins in all significantly enriched pathways and networks. This analysis does not provide the means to statistically evaluate the significance of miRNA addition to a given pathway or network; however, this approach provides evidence of an epigenetic component of the liver response to HFr diet.

### Identification of pathway and network genes previously associated with NASH/NAFLD related traits

The following search terms, with all variation of names in the GWAS catalog, were used to query the current GWAS catalog (74): alkaline phosphatase, aspartate aminotransferase, body mass index, body weight, fasting blood glucose, fasting blood insulin, fat body mass, fatty acid, glucose, HbA1c, HDL cholesterol change, insulin, insulin resistance, insulin sensitivity, LDL cholesterol change, lipid, liver fat, liver disease biomarker, liver fibrosis, low density lipoprotein cholesterol, non-alcoholic fatty liver disease, non-alcoholic steatohepatitis, obesity, omega-3 polyunsaturated fatty acid, omega-6 polyunsaturated fatty acid, total cholesterol, triglyceride, type II diabetes mellitus, very low density lipoprotein cholesterol. Genes with associations, based on the GWAS catalog, to any of these traits were compared to the list of all differentially expressed miRNAs, genes and proteins from our transcriptomic and proteomic datasets, and compared with the genes in proteins in multi-omic significant networks and pathways.

## LIST OF ABBREVIATIONS

HFr: high fructose
PPARA: peroxisome proliferator activated receptor alpha
DHA: docosahexaenoic acid
NHP: nonhuman primates
NASH: nonalcoholic steatohepatitis
NAFLD: nonalcoholic fatty liver disease
GEO: Gene Expression Omnibus
H-MCR: hierarchical multivariate curve resolution
MeOX: methoxyamine hydrochloride
MSTFA: N-methyl-N-trimethylsilyl-trifluoroacetamide
EI: electron impact
RI: retention indices
AMDIS: Automated Mass Spectral Deconvolution and Identification System
NIST: National Institute of Standards and Technology
LC-TOFMS: Liquid Chromatography-Time of Flight Mass Spectrometry
HMDB: Human Metabolome Database
PCA: principle component analysis
VIP: variable importance in the projection
IPA: Ingenuity Pathway Analysis

## DECLARATIONS

### Ethics approval and consent to participate

All experimental procedures involving vervet monkeys (*Chlorocebus sabaeus*) were approved and complied with the guidelines of the Institutional Animal Care and Use Committee of Wake Forest University Health Sciences, which is an AALAC accredited facility. Procedures were performed by a board-certified veterinarian employed by Wake Forest University Health Sciences.

### Consent for publication

All authors have reviewed the manuscript and consent for publication.

### Availability of data and materials

RNA Seq, proteomic, and metabolomic data are available in Additional files. Raw RNA Seq data are available through NCBI GEO Series accession number GSE176576 and small RNA Seq data are available through GEO accession number GSE178269.

To review GEO accession GSE178269 go to:

https://urldefense.com/v3/__https://www.ncbi.nlm.nih.gov/geo/query/acc.cgi?acc=GSE178269__;!!GA8Xfdg!hpk17gHazQbLJ2Ux3IeSzs9VDjsSSqkQ9zsGAAIMuyNtd_NsH2pPRYGKi1hk9Jw$

The following secure token has been created to allow review of record GSE178269 while it remains in private status: yryzskgcjrqfxav

To review GEO accession GSE176576 go to:

https://urldefense.com/v3/__https://www.ncbi.nlm.nih.gov/geo/query/acc.cgi?acc=GSE176576__;!!GA8Xfdg!iPSaGb7UXXtZbc2VRI0Nj1cw9VE7rn_cFK62irhMe4UbjQs4vcXTLI31lSgLr38$

The following secure token has been created to allow review of record GSE178269 while it remains in private status: knsrqcaodtoxjgz

### Authors’ contributions

LAC, KK, and MO conceived the project. JPG, AJ, PR, GMK, LAC, JC, ZH, EQ, VD, and MO contributed to data generation and analyses. All authors read and approved the final manuscript.

## Acknowledgements

We thank Biswapriya Misra for contributing to generation of metabolomics data.

## Supplementary Information

**Additional file 1**

Pathway Summary for genes, proteins, metabolites, combined genes and proteins, combined genes, proteins, and metabolites, and combined genes, proteins, metabolites, and miRNAs.

**Additional file 2**

Network Summary for genes, proteins, metabolites, combined genes and proteins, combined genes, proteins, and metabolites, and combined genes, proteins, metabolites, and miRNAs.

**Additional file 3**

Gene List: Genes passing quality filters with ratios and p-values for HFr versus CON.

**Additional file 4**

Gene Pathways: Enrichment analysis of genes with p-value < 0.05.

**Additional file 5**

Gene Networks: Enrichment analysis of genes with p-value < 0.05.

**Additional file 6**

Protein List: Proteins passing quality filters with ratios and p-values for HFr versus CON.

**Additional file 7**

Protein Pathways: Enrichment analysis of proteins with p-value < 0.05.

**Additional file 8**

Protein Networks: Enrichment analysis of proteins with p-value < 0.05.

**Additional file 9**

Common Genes Proteins: List of common genes and proteins from Venney merge for all genes and proteins passing quality filters and for all differentially expressed genes and proteins.

**Additional file 10**

Metabolite List: Metabolites passing quality filters with ratios and p-values for HFr versus CON.

**Additional file 11**

Metabolite Pathways: Enrichment analysis of metabolites with p-value < 0.05.

**Additional file 12**

Metabolite Networks: Enrichment analysis of metabolites with p-value < 0.05.

**Additional file 13**

Gene & Pro Pathways: Enrichment analysis combining genes and proteins with p-value < 0.05.

**Additional file 14**

Gene & Pro Networks: Enrichment analysis combining genes and proteins with p-value < 0.05.

**Additional file 15**

Gene Pro Met Pathways: Enrichment analysis combining genes, proteins, and metabolites with p-value < 0.05.

**Additional file 16**

Gene Pro Met Networks: Enrichment analysis combining genes, proteins, and metabolites with p-value < 0.05.

**Additional file 17**

miRNA List: miRNAs passing quality filters and p-values < 0.05 for HFr versus CON.

**Additional file 18**

Gene-Pro with miRNA pairs: miRNA pairing with target genes and proteins either highly predicted or experimentally validated for differentially expressed miRNAs, genes and proteins for HFr versus CON (p-value <0.05).

**Additional file 19**

Diff Gene Pro GWAS: List of GWAS hits of differentially expressed genes and proteins for HFr versus CON.

